# A Bayesian Framework for Multiple Trait Colocalization from Summary Association Statistics

**DOI:** 10.1101/155481

**Authors:** Claudia Giambartolomei, Jimmy Zhenli Liu, Wen Zhang, Mads Hauberg, Huwenbo Shi, James Boocock, Joe Pickrell, Andrew E. Jaffe, the CommonMind Consortium, Bogdan Pasaniuc, Panos Roussos

## Abstract

**Motivation:** Most genetic variants implicated in complex diseases by genome-wide association studies (GWAS) are non-coding, making it challenging to understand the causative genes involved in disease. Integrating external information such as quantitative trait locus (QTL) mapping of molecular traits (e.g., expression, methylation) is a powerful approach to identify the subset of GWAS signals explained by regulatory effects. In particular, expression QTLs (eQTLs) help pinpoint the responsible gene among the GWAS regions that harbor many genes, while methylation QTLs (mQTLs) help identify the epigenetic mechanisms that impact gene expression which in turn affect disease risk. In this work we propose **m**ultiple-trait-c**oloc** (*moloc*), a Bayesian statistical framework that integrates GWAS summary data with multiple molecular QTL data to identify regulatory effects at GWAS risk loci.

**Results:** We applied *moloc* to schizophrenia (SCZ) and eQTL/mQTL data derived from human brain tissue and identified 52 candidate genes that influence SCZ through methylation. Our method can be applied to any GWAS and relevant functional data to help prioritize disease associated genes.

**Availability:** *moloc* is available for download as an R package (https://github.com/clagiamba/moloc). We also developed a web site to visualize the biological findings (icahn.mssm.edu/moloc). The browser allows searches by gene, methylation probe, and scenario of interest.

**Supplementary information:** Supplementary data are available at *Bioinformatics* online.

## 1 Introduction

Genome-wide association studies (GWAS) have successfully identified thousands of genetic variants associated with complex diseases (Visscher *et al.*, 2012). However, the majority of the discovered associations point to non-coding regions, making it difficult to identify the causal genes and the mechanism by which risk variants mediate disease susceptibility. A potential approach to explore the mechanism of risk non-coding variants is through integration with datasets that measure the association of molecular phenotypes such as gene expression (expression QTL or eQTL) and DNA methylation (methylation QTL or mQTL). The observation that the same variant is driving the association signal in GWAS, and also affecting expression at a near-by gene and methylation site, could indicate a putative dis-ease mechanism. Analyzing two datasets jointly has been a successful strategy to identify shared genetic variants that affect different molecular processes, in particular eQTL and GWAS (Fromer *et al.*, 2016; Zhu *et al.*, 2016; Hauberg *et al.*, 2017; Gusev *et al.)* and mQTL and GWAS integration (Jaffe *et al.*, 2016; Hannon, Dempster, *et al.*, 2016; Hannon, Spiers, *et al.*, 2016; Hannon *et al.*, 2017). All these previous efforts have focused on pairwise dataset integration (e.g. eQTL and GWAS or mQTL and GWAS).

To our knowledge, a statistical approach to integrate more than two datasets with information on genetic associations is lacking. Therefore, we developed *multiple*-trait-*coloc* (*moloc*), a statistical method to quantify the evidence in support of a common causal variant at a particular risk region across multiple traits. Our approach is a multi-trait extension of our previously developed two-trait model described in *coloc* (Claudia Giambartolomei *et al.*, 2014). This method can be used to compare association signals for multiple phenotypes (molecular or complex disease traits), using summary-level information from genetic association datasets.

To illustrate the advantage of a joint analysis in real data, we applied *moloc* to schizophrenia (SCZ), a complex polygenic psychiatric disorder, using summary statistics from the most recent and largest GWAS by the Psychiatric Genomics Consortium (Schizophrenia Working Group of the Psychiatric Genomics Consortium, 2014), which reported association for 108 independent genomic loci. eQTL data were derived from the CommonMind Consortium (Fromer *et al.*, 2016), which generated the largest eQTL dataset in the dorsolateral prefrontal cortex (DLPFC) from SCZ cases and control subjects (N=467). Finally, we leveraged mQTL data that were previously generated in human DLPFC tissue (N=121) to investigate epigenetic variation in SCZ (Jaffe *et al.*, 2016). Integration of multiple phenotypes helps better characterize the genes predisposing to complex diseases such as SCZ.

## 2 Methods

### 2.1 Method Description

We introduce *moloc* to detect colocalization among any number of traits in a specific locus. The input of the model is the set of summary statistics derived from three (or more) traits measured in distinct datasets of unrelated individuals. In this manuscript, we refer to traits (e.g. complex trait, gene expression and DNA methylation) as a synonymous to datasets containing the information on genetic associations (e.g. GWAS, eQTL and mQTL).

We define a genomic region containing Q variants, for example a *cis* region around expression or methylation probe. We are interested in a situation where summary statistics (effect size estimates and standard errors) are available for all datasets in the genomic region. We first derive our model using three traits, then generalize to any number of traits. If we consider colocalization of three traits (GWAS, eQTL and mQTL), under a maximum of a single causal variant per trait, there can be up to three causal variants and 15 possible scenarios summarizing how the variants are shared among the traits. Each hypothesis can be represented by a set of index sets according to which of the traits each SNP is associated with (all hypotheses are listed in **Table S1**).

To illustrate our notation, consider a region with 8 SNP. For simplicity, we denote GWAS as G, eQTL as E, and mQTL as M. Four examples of configurations are shown in Figure 1. The “.” in the subscript denotes scenarios supporting different causal variants. For instance, GE summarizes the scenario for one causal variant shared between traits GWAS and eQTL (Figure 1 - Right plot top panel); GE.M summarizes the scenario with one causal variant for traits GWAS and eQTL, and a different causal variant for trait mQTL (Figure 1 - Left plot bottom panel).

**Figure 1.**
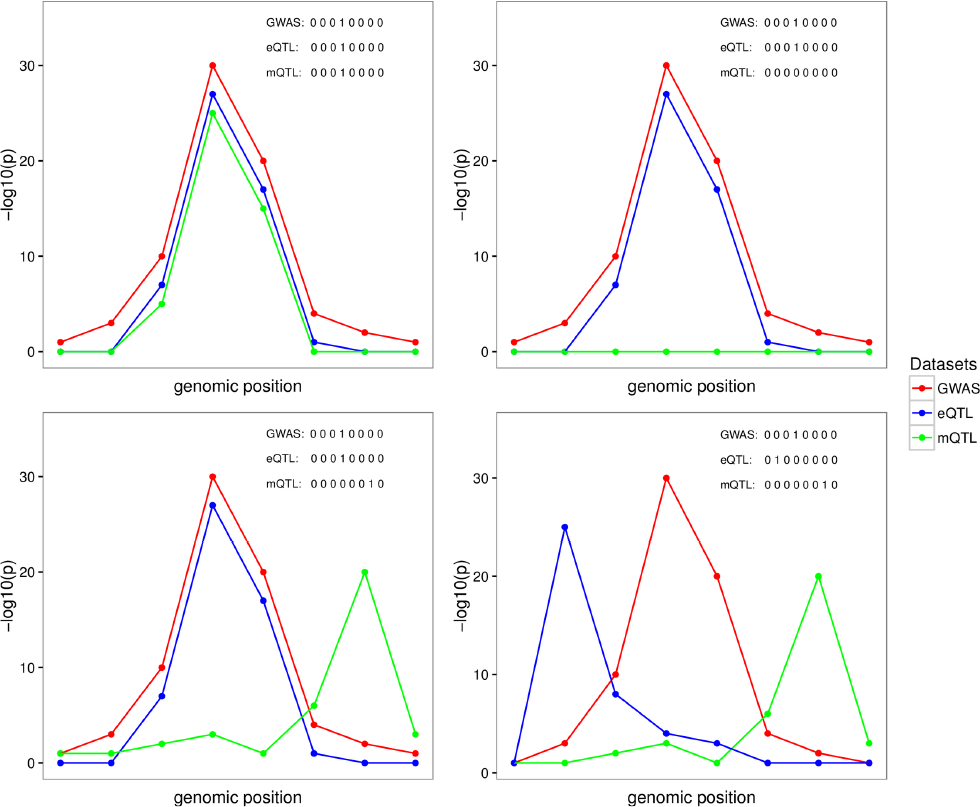
Graphical representation of four possible configurations at a locus with 8 SNPs in common across three traits. The traits are labeled as G, E, M representing GWAS (G), eQTL (E), and mQTL (M) datasets, respectively. Each plot represents one possible configuration, which is a possible combination of 3 sets of binary vectors indicating whether the variant is associated with the selected trait. Left plot top panel (GEM scenario): points to one causal variant behind all of the associations; Right plot top panel (GE scenario): represent the scenario with the same causal variant behind the GE and no association or lack of power for the M association; Left plot bottom panel (GE.M scenario): represents the case with two causal variants, one shared by the G and E, and a different causal variant for M; Right plot bottom panel (G.E.M. scenario): represents the case of three distinct causal variants behind each of the datasets considered.

Our approach computes the evidence supporting the 15 possible scenarios (H0…H14), of sharing of SNPs among traits in the given genomic region. We first compute the posterior probability of any of the 15 configurations by weighting the likelihood of the data D given a configuration S, P(D|S), by the prior probability of a configuration, P(S) (described below). We can reformulate the posterior probability for each hypothesis as a ratio by dividing each by the baseline likelihood supporting the first model of no association with any trait H0. The probability of the data for hypothesis h is then the sum over all configurations Sh, which are consistent with the given hypothesis:

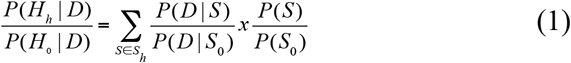

where, P(D|S)/P(D|S0) is the Bayes Factor for each configuration compared to the baseline configuration of no association with any trait S0, P(S)/P(S0) is the prior odds of a configuration compared with the baseline configuration S0, and the sum is over Sh, the set of configurations supporting hypothesis H0 to H14. Similar to pairwise colocalization (Claudia Giambartolomei *et al.*, 2014) we then estimate the evidence in support of different scenarios in a given genomic region using the posterior probability supporting hypothesis h among H possible hypothesis, computed from:

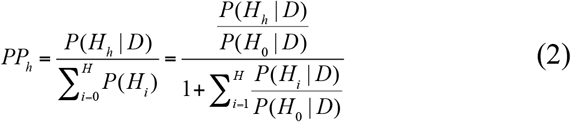

Therefore, in our application, the method outputs 15 posterior probabilities. We are most interested in the scenarios supporting a shared causal variant for two and three traits.

We make three important assumptions in *moloc*, the same that are made in our previous *coloc* methodology. Firstly, that the causal variant is included in the set of **Q** common variants, either directly typed or well imputed. If the causal SNP is not present, the power to detect a common variant will be reduced depending on the linkage disequilibrium (LD) between other SNPs included in the model and the causal SNP. Secondly, we assume at most one causal variant is present for each trait per locus. In the presence of multiple causal variants per trait, this method is not able to identify colocalization between additional association signals independent from the primary one. Thirdly, as we do not explicitly model LD between SNPs, we assume the samples are drawn from the same ethnic population and therefore have identical allele frequencies and patterns of LD.

### 2.2 Bayes Factor of a SNP with one trait

We start by computing a Bayes Factor for each SNP and each trait (i.e GWAS, eQTL, mQTL). We assume a simple linear regression model to relate the phenotypes or a log-odds generalized linear model for the case-control dataset, and the genotypes. Using the Wakefield Approximate Bayes factors (Wakefield, 2009) (WABF), only the variance and effect estimates from regression analysis are needed, as shown below and previously described (Pickrell *et al.*, 2016; Claudia Giambartolomei *et al.*,2014):

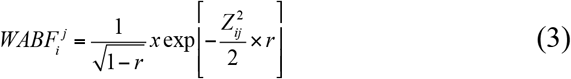

where Z_ij_ = 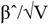 is the usual Z statistic and the shrinkage factor r is the ratio of the variance of the prior and total variance (r = W/(V + W)).

The WABF requires specifying the variance W of the normal prior. In the *moloc* method we set W to 0.15 for a continuous trait and 0.2 for the variance of the log-odds ratio parameter, as previously described (Claudia Giambartolomei *et al.*, 2014). Another possibility is to average over Bayes factors computed with W = 0.01, W = 0.1, and W = 0.5 (Pickrell *et al.*, 2016). We provide this as an option that can be specified by the user. If the variance of the estimated effect size V is not provided, it can be approximated using the allele frequency of the variant f, the sample size N (and the case control ratio s for binary outcome) (Claudia Giambartolomei *et al.*, 2014):

### 2.3 Bayes factor of a SNP across more than one trait

To compute the BF where a SNP i associates with more than one trait, we use:

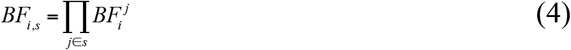

Where **s** is the set of trait indices for which SNP i is associated with. Note that the computations under >1 trait multiply the individual Bayes Factors together. This is equivalent to the Bayes Factor under the maximum heterogeneity model used in Wen and Stephens (Wen and Stephens, 2011). Two key assumptions are necessary for the following computations. Firstly, that the traits are measured in unrelated individuals. The datasets we used in the current analysis does not contain overlapping individuals, however we provide the code to adjust for this. Secondly that the effect sizes for the two traits are independent (Claudia Giambartolomei *et al.*, 2014).

### 2.4 Prior probabilities that SNP i associates with traits indexed in s

The prior probability that SNP i associates with all traits indexed in a set in our three trait model is: π_φ_ SNP i associates with no trait, with one trait, pairs or traits or all traits π{1,2,3} such that they sum to 1: π_φ_ + π_{1}_ + π_{2}_ + π_{3}_ + π_{1,2}_ + π_{1,3}_ + π_{2,3}_ + π_{1,2,3}_ = 1.

### 2.5 Simplified model under the assumption that all SNPs have the same prior

The probability of the data under each hypothesis can be computed by summing the probability of all the causal configurations consistent with a particular hypothesis, weighted by the prior probabilities (**Equation 1**). In our model, P(S) is the prior probability under any one of the 15 hypotheses. We can define these priors from the prior probability that a SNP *i* associates with traits indexed in *π_s_* (section above). Additionally, since the prior probability P(S) of any one configuration in the different sets do not vary across SNPs that belongs to the same set *Sh*, we can multiply the likelihoods by one common prior supporting the different hypothesis (Claudia Giambartolomei *et al.*, 2014). In this framework, we can see that P(S) depends on a ratio of *π_s_* and on **Q**, the number of SNPs in the region (**Text S1**), Therefore, across a set of *j* traits {1,2,3…,}, we compute the probability of the data supporting hypothesis h, where one SNP is associated with j traits as:

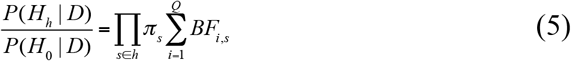

where BF_i,s_ are the Bayes factor of a SNP across traits indexed in s (**Equation 4**), *π* are the prior probabilities that SNP i is the causal SNP under a specific model.

The probability of the data where there are more than one independent associations among the j traits (i.e. |h| > 1) can be derived from the pre-computed probability of the data where there is one association among the j traits (i.e. |h| = 1, **Text S1**):

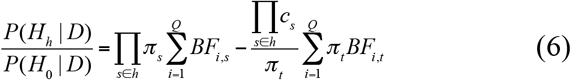

### 2.6 Prior Probabilities of each hypothesis

In practice, we collapsed the prior probabilities to a smaller set for each kind of configuration. We set the prior probability that a SNP is causal in each trait to be identical (π_{1}_ = π_{2}_ = π_{3}_) and refer to this a p1. We also set the prior probability that is associated with two traits to be identical (π_{1,2}_, π_{2,3}_, π_{1,3}_) and refer to this as p2. We refer to the prior probability that SNP i the causal for all traits (π_{1,2,3}_) as p3.

### 2.7 Moloc analysis

The GWAS, eQTL, mQTL datasets were filtered by minor allele frequency greater than 5% and had individually been filtered by imputation quality (**Text S1**). The Major Histocompatibility (MHC) region (chr 6: 25 Mb - 35 Mb) was excluded from all co-localization analyses due to the extensive linkage disequilibrium and complexity of the associations. We applied a genic-centric approach, defined *cis*-regions based on a 50kb upstream/downstream from the start/end of each gene, since our goal is to link risk variants with changes in gene expression. We evaluated all methylation probes overlapping the *cis*-region. The number of *cis*-regions/methylation pairs is higher than the count of genes because, on average, there are more than one methylation sites per gene. Common SNPs were evaluated in the colocalization analysis for each gene, and each methylation probe, and GWAS. In total, 12,003 *cis*-regions and 481,995 unique *cis*-regions/methylation probes were tested. Genomic regions were analyzed only if greater than 50 SNPs were in common between all the datasets. Across all of the analyses, a posterior probability equal to, or greater than, 80% for each configuration was considered evidence of colocalization.

In order to compare colocalization of two trait analyses with three traits, we applied our previously developed method [*coloc* (Claudia Giambartolomei *et al.*, 2014)]. Effect sizes and variances were used as opposed to p-values, as this strategy achieves greater accuracy when working with imputed data (Claudia Giambartolomei *et al*., 2014).

### 2.8 Simulations

We simulated genotypes from sampling with replacement among haplotypes of SNPs with a minor allele frequency of at least 5% found in the phased 1000 Genomes Project within 49 genomic regions that have been associated with type 1 diabetes (T1D) susceptibility loci (excluding the major histocompatibility complex (MHC) as previously described (Wallace, 2013). These represent a range of region sizes and genomic topography that reflect typical GWAS hits in a complex trait. For each trait, two, or three “causal variants” were selected at random. We have simulated continuous traits, and assume that causal effects follow a multivariate Gaussian distribution, with each causal variant explaining 0.01 variance of the trait in the GWAS data, and 0.1 in the eQTL and mQTL datasets. Note that colocalisation testing may be applied equally to quantitative data (using linear regression), and to case control data (using logistic regression). For the null scenario, the causal variants explain zero variance of the traits. To quantify false positive rates on a large number of tests, we simulated the null 500,000 times. We simulated the 15 possible scenarios with different sharing patterns between the GWAS, eQTL, and mQTL datasets. We used sample sizes of 82,315, 467, and 121 individuals to reflect our true sample sizes. We also used different combinations of sample sizes to explore power to detect the correct hypothesis.

We estimated the number of false positives within each simulated scenario, by counting the proportion of simulations under the null that passed a posterior probability supporting each of the 14 hypothesis at a particular threshold (PPA>=threshold). We also report the false positives using the sum of the posteriors (PPA.ab + PPA.ab.c + PPA.abc). The false positive rate is the number of false positives over 1,000 simulations. We repeated this procedure using 500,000 simulations under our true sample sizes.

We next sought to compare the misclassification rates, and power to detect the correct hypothesis. To compute the number of misclassified calls within each simulated scenario, we counted the proportion of simulations that passed a posterior probability supporting a different hypothesis from the one simulated at a particular threshold (PPA>=threshold). We estimated power to distinguish a particular hypothesis from the others by counting the proportion of correctly identified simulations at a particular threshold (PPA(true)>=threshold). Since in most cases the causal variant will not be included in the panel, we repeated simulations after removing the causal variant.

To explore the effect of linkage disequilibrium (LD) on estimated posterior probability we first computed an LD score for each SNP in the region, defined as the sum of the squared correlation between a SNP and all the SNPs in the region. To assess the degree of LD at a locus we took the average of these scores. All analyses were conducted in R.

## 3 Results

### 3.1 Sample Size Requirements

In a first set of simulations we explored false positive rates (**Figure S1**) and the posterior probability under different sample sizes (**Figures S2, S3**). False positive rates are below 0.05 even if a threshold of 0.3 for posteriors is used, and where the causal variant is masked (**Figures S1**). Figure 2A illustrates the posterior probability distribution across our three scenarios of interest: GWAS and eQTL, alone or together with mQTL. With a GWAS sample size of 10,000 and eQTL and mQTL sample sizes of 300, the method provides reliable evidence to detect a shared causal variant behind the GWAS and another trait (median posterior probability of any hypothesis >50%). The posterior across all of the possible scenarios is illustrated in **Figure S2**. Although in this paper we analyze GWAS, eQTL and mQTL, our method can be applied to any combinations of complex disease and molecular traits, including 2 GWAS traits and an eQTL dataset. We explored the minimum sample size required when analyzing two GWAS datasets (termed G1, G2) and one eQTL (E) (**Figure S3**). The method provides reliable evidence for all hypotheses when the two GWAS sample sizes are 10,000 and eQTL sample size reaches 300. We then explored misclassification rates (**Tables S2, S3, S4**). When samples are 10,000 for GWAS and greater than 300 for eQTL and mQTL, misclassification rates for detecting our hypotheses of interests at 80% threshold are all below 0.05 (**Table S2**). Where the causal variant is masked, sample sizes also need to reach 10,000 for GWAS and greater than 300 for eQTL and mQTL, for misclassification rates to be below 0.05 (**Table S3**). Given the small sample size for the mQTL data, the method has trouble detecting a different causal variant for the mQTL dataset (**Table S4**). For example, evidence pointing to two different causal variants between GWAS and eQTL could be generated by the presence of three causal variants in reality, but the causal variant for mQTL remains undetected. For this reason, we focused on cases with shared casual variants between GWAS, eQTL, with or without mQTL.

**Figure 2.**
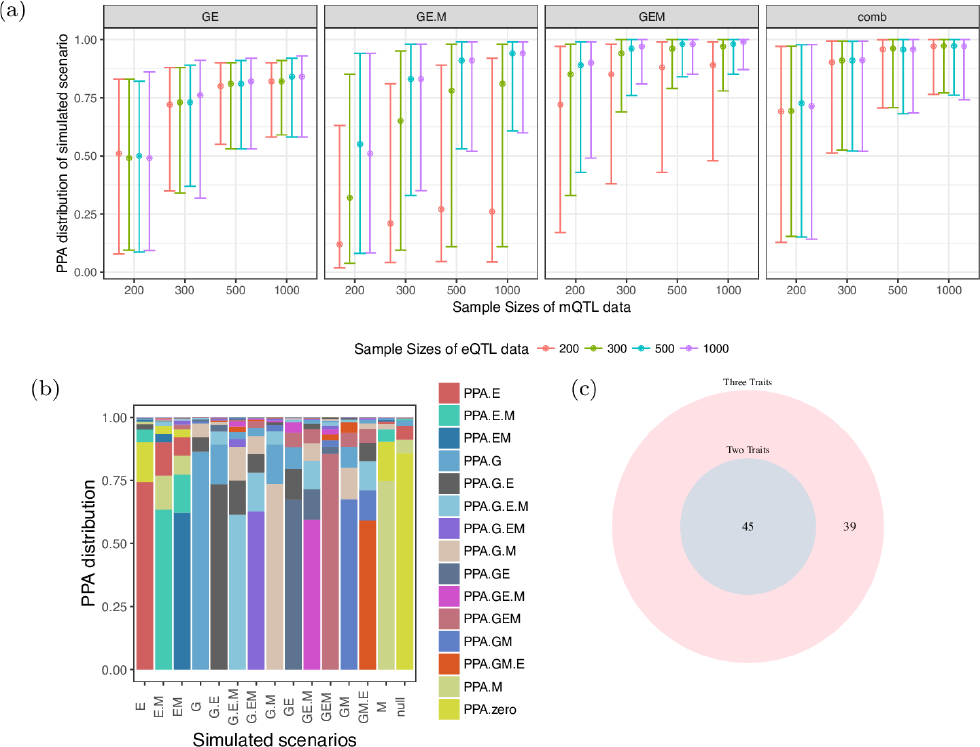
Results from simulations under colocalization/noncolocalization scenarios (A-B), and results from real data application (C). A. Simulations under different sample sizes for all scenarios in moloc of three traits (GWAS, eQTL, mQTL). The y axis shows the median, 10% and 90% quantile of the distribution of posterior probabilities (“PPA”), which supports each of our scenarios of interest. Combined scenarios include gene-methylation pairs or genes that reach a posterior probability of GEM >= 80%, or + GE.M>= 80%, or GE>= 80%. All cases include 10,000 individuals in the GWAS dataset. The variance explained by the trait was set to 0.01 for GWAS (1%), and to 0.1 (10%) for the eQTL and mQTL. B. Posterior probabilities from simulations using a sample size of 10,000 individuals for GWAS trait (denoted as G), G), 300 for eQTL trait (denoted as E), and 300 for mQTL trait (denoted as M). X-axis shows all 15 simulated scenarios, e.g. G.E.M, three different causal variants for each of the three traits. Y-axis shows the distribution of posterior probabilities under the simulated scenario. The height of the bar represents the mean of the PPA for each configuration across simulations. C. Venn diagram comparing number of colocalization of two traits (coloc PPA >=80%) with three traits (moloc PPA GE + GE.M + GEM) in simulations with one causal variant shared between all the three traits (GEM). Results include 887 out of 1,000 simulations passing 80% threshold for colocalization.

It is instructive to observe where evidence for other hypotheses is distributed. Figure 2B illustrates the accuracy of our approach under different scenarios where two or three causal variants are shared. For example, under simulations of one shared variant for GWAS and eQTL and a second variant for mQTL (GE.M), on average 60% of the evidence points to the simulated scenario, while 12% point to GE, 12% to G.E.M and 7.2% to GEM.

### 3.2 Choice of priors

The method requires the definition of prior probabilities for the association of a SNP with one (p1), two (p2), or three traits (p3). We set the prior probability that a variant is associated with one trait as 1 x 10^−4^ for GWAS, eQTL and mQTL, assuming that each genetic variant is equally likely a priori to affect gene expression or methylation or disease. This estimate has been suggested in the literature for GWAS (Stephens and Balding, 2009) and used in similar methods (Hormozdiari *et al.*, 2016). We set the priors p2 = 1 x 10^−6^, p3 = 1 x 10^−7^ based on sensitivity and exploratory analysis of genome-wide enrichment of GWAS risk variants in eQTLs and mQTLs. In **Figure S4**, we find eQTLs and mQTLs to be similarly enriched in GWAS, justifying our choice of the same prior probability of association across the two traits. These values are also suggested by a crude approximation of p2 and p3 from the common genome-wide significant SNPs across the three datasets.

We performed sensitivity analyses using different priors. Specifically, we fixed p1 to 1×10^−4^ and tested a range of priors for p2 and p3 from 1×10^−5^ to 1×10^−8^, with increasing difference between p1, p2 and p3. We used a form of internal empirical calibration to compare our prior and posterior expectations. We find that the posterior expectation of colocalization most closely resembled the prior expectation under our choice of priors (**Table S5**). We note that our R package implementation allows users to specify a different set of priors. Additionally, we could vary the prior probability of a SNP to be causal based on features of interest, using estimates of the prior probability of a SNP to be causal given specific annotations (Kichaev *et al.*, 2014; Pickrell, 2014; Li and Kellis, 2016; Chung *et al.*, 2014). We demonstrate this utility by applying *fGWAS* (Pickrell, 2014) to the SCZ dataset, together with different chromatin marks measured in the dorsolateral prefrontal cortex as profiled by the Roadmap Epigenomics Consortium (see Text S1).

### 3.3 Co-localization of eQTL, mQTL and risk for Schizophrenia

We applied our method to SCZ GWAS using eQTLs derived from 467 samples and mQTL from 121 individuals (Text S1). Our aim is to identify the genes important for disease through colocalization of GWAS variants with changes in gene expression and DNA methylation. We analyzed associations genome-wide, and report results both across previously identified GWAS loci, and across potentially novel loci. While we consider all 15 possible scenarios of colocalization, here we focus on gene discovery due to higher power in our eQTL dataset, by considering the combined probabilities of cases where the same variant is shared across all three traits GWAS, eQTLs and mQTLs (GEM > 0.8) or scenarios where SCZ risk loci are shared with eQTL only (GE > 0.8 or GE.M > 0.8) (Table 1 and **Table S1**). We identified 1,053 cis-regions/methylation pairs with posterior probability above 0.8 that are associated with all three traits (GEM), or eQTLs alone (GE or GE.M). These biologically relevant scenarios affect overall 84 unique genes and include 39 genes that fall within the previously identified SCZ LD blocks (**Table S6**) and 45 potentially novel genes outside of these regions (**Table S7**). Fifty-two out of the 84 candidate genes influence SCZ, gene expression and methylation (GEM>=0.8). One possible scenario is that the variants in these genes could be influencing the risk of SCZ through methylation, although other potential interpretations such as pleiotropy should be considered.

**Table 1.**
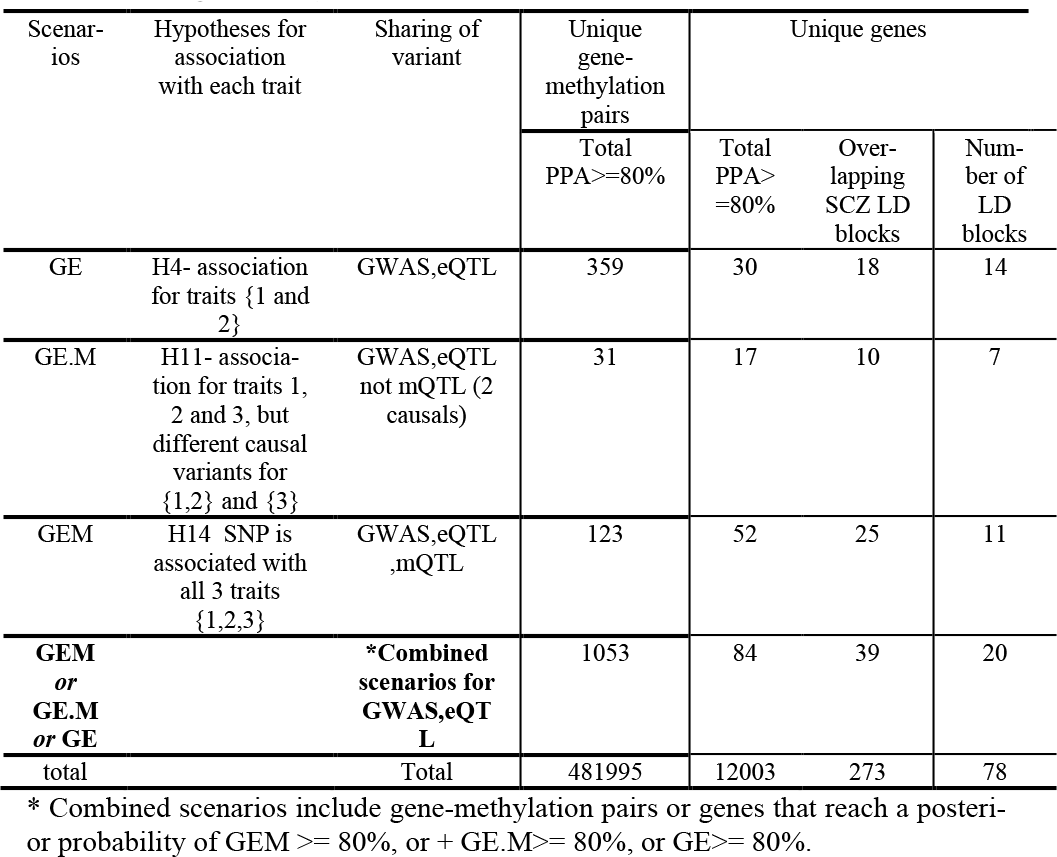
Number of genes with evidence of colocalization (PPA>=0.8) under each scenario.

### 3.4 Addition of a third trait increases gene discovery

We examined whether *moloc* with 3 traits enhance power for GWAS and eQTL colocalization compared to using 2 traits. In simulations to compare *coloc* and *moloc* under one causal variant and our true sample sizes for all three datasets, we observe a fold increase of 1.5 for gene discovery using *moloc* versus *coloc*. *Moloc* with three traits recovers all the genes discovered using coloc with eQTL and mQTL, and additional genes from the inclusion of the third layer. In our real data, colocalization analysis of only GWAS and eQTL traits identified 45 genes with a posterior probability, PP4 in *coloc*, of >= 0.8. The 39 additional genes that were found by adding methylation include genes such as *CALN1*, a neuronal transcript associated with abnormalities in sensorimotor gating in humans (Roussos *et al.*, 2016), that would have been missed by only GWAS and eQTL colocalization.

### 3.5 Loci overlapping reported SCZ LD blocks

Psychiatric Genomics Consortium (PGC) identified 108 independent loci and annotated LD blocks around these, 104 of which are within non-HLA, autosomal regions of the genome (Schizophrenia Working Group of the Psychiatric Genomics Consortium, 2014). In Table 1 and **Table S1** we report the number of identified gene-methylation pairs and unique genes under each scenario that overlap the SCZ-associated LD blocks. Out of the 78 SCZ-associated LD blocks we examined in our analysis, we found colocalizations in 20 of them. We note that each LD block can cover multiple genes that co-localize with the GWAS signal; in fact within a block, there are, on average, 2.4 unique genes that reach evidence of sharing the same causal variant with a GWAS signal. **Figure S5A** illustrates the average distribution of the posteriors across these regions. Cumulatively, 12% of the evidence points to shared variation with an eQTL (GE, GE.M and GEM). The majority of the evidence within these regions (64%) did not reach support for shared variation across the three traits, with 20% not reaching evidence for association with any traits, and 44% with only one of the three traits (36% with GWAS; 6% with eQTL, 2% with mQTLs). The lack of evidence in these regions could be addressed with greater sample sizes. **Figure S5B** shows the evidence for colocalization of GWAS with eQTL or mQTL across the 39 candidate genes. We provide illustrative examples of SCZ association with expression and DNA methylation in the *FURIN* locus (**Figures S6** and **S7**).

### 3.6 Potentially novel SCZ loci

We found 45 unique genes that have a high posterior for SCZ and eQTL, but fall in regions not previously identified to be associated with SCZ (at p-value of 5 × 10^−8^). All genes were far from a SCZ LD block (more than 150kb, **Table S7**), and contained SNPs with p-values for association with SCZ ranging from 10^−4^ to 10^−8^. These genes will likely be identified using just the GWAS signal if the sample size is increased. *KCNN3* is among these genes which encodes an integral membrane protein that forms a voltage-independent calcium-activated channel. It regulates neuronal excitability by contributing to the slow component of synaptic after hyperpolarization (Deignan *et al.*, 2012). A plot of the associations with the three datasets within this locus is shown in **Figure S6B**.

### 3.7 Comparison with previous findings

We compare our gene discovery results to previous studies that assess GWAS-eQTL (Fromer *et al.*, 2016; Zhu *et al.*, 2016; Hauberg *et al.*,2017; Gusev *et al.)* or GWAS-mQTL (Hannon, Dempster, *et al.*, 2016; Gusev *et al.;* Hannon, Spiers, *et al.*, 2016; Hannon *et al.*, 2017) colocalization using the same or similar datasets (**Table S9** and **Figure S8**). A substantial proportion of genes detected in our study (range 44%-85%, pairwise hypergeometric p-value <0.01) was validated with four studies (Fromer *et al.*, 2016; Zhu *et al.*, 2016; Hauberg *et al.*, 2017; Gusev *et al.)* that used eQTL and GWAS integration to prioritize genes important for schizophrenia. Several studies have also linked methylation data with schizophrenia (Hannon, Dempster, *et al.*, 2016; Hannon, Spiers, *et al.*,2016; Hannon *et al.*, 2017). Two recent studies (Hannon, Dempster, *et al.*, 2016; Hannon *et al.*, 2017) used blood mQTL data from 639 samples and identified colocalization of SCZ loci with 32 and 200 methylation probes by applying *coloc* and *SMR*, respectively. A proportion of SCZ-mQTL colocalization was validated in our study (*coloc:* 46%; *SMR:*18%, **Table S9**). Overlap between our analysis in brain and these analyses point to shared mechanisms in blood and brain. Another study (Hannon, Spiers, *et al.*, 2016) used mQTL data from 166 fetal brain samples and identified 297 methylation probes important for schizophrenia. We analyzed 184 of those and found evidence for 13 probes. We note that our methylation data did not included fetal brain samples. Finally, a recent study (Gusev *et al.*) identified 44 genes involved in schizophrenia through TWAS, followed by integration with chromatin data in blood that resulted in 11 genes associated with GWAS, eQTL and epigenome QTL. We analyzed 8 out of the 11 associations and confirmed 6 of these genes that, in our study, influence SCZ through eQTL and mQTL.

### 3.8 Association of gene expression with methylation

We examined the association of DNA methylation and gene expression as a function of distance from transcription start site. We explored direction of effects of methylation and expression, for gene expression and DNA methylation that colocalize (PPA.GEM + PPA.EM + PPA.G.EM >= 0.8). This approach has the advantage of linking changes in methylation with specific transcripts, avoiding the issues of arbitrary annotating CpG methylation sites to the nearby genes. Overall, we tested 1,947 DNA methylation and gene expression pairwise interactions and found a significant negative correlation between the effect sizes of methylation and expression in the proximity of the transcription start site (**Figure S9**, p-value:<2.2×l0^−16^). **Table S8** provides a list of methylation and gene expression pairwise interactions in human brain tissue for DNA methylation probes that are proximal to the transcription start site (20kb upstream to 2kb downstream of transcription start site).

## 4 Discussion

In this paper, we propose a statistical method for integrating genetic data from molecular quantitative trait loci (QTL) mapping into genome-wide genetic association analysis of complex traits. The proposed approach requires only summary-level statistics and provides evidence of colocalization of their association signals. To our knowledge, a method integrating more than two traits is lacking. In contrast to other methods that attempt to estimate the true genetic correlation between traits such as LD score regression (Bulik-Sullivan *et al.*, 2015) and TWAS (Gusev *et al*.), *moloc* focuses on genes that are detectable from the datasets at hand. Thus, if the studies are underpowered, most of the evidence will lie in the null scenarios. We note that our model is the same as *gwas-pw* in Pickrell et al. (Pickrell *et al.*, 2016) under specific settings. Precisely, *gwas-pw* averages over Bayes factors computed with W = 0.01, W = 0.1, and W = 0.5 (Methods). We provide this as an option that can be specified by the user. Additionally, *gwas-pw* estimates the prior parameters genome-wide using a maximization procedure. However, we note that, unlike *gwas-pw* that focuses on genome-wide estimation across pairs of traits, our approach focuses on one locus at a time with multiple traits. Our method is complementary to methods quantifying local genetic covariance across traits. The aim of this method is to identify cases where the same causal variant is shared between the traits. We argue that it is valuable to identify cases where the same signal is influencing multiple traits, for example when studying application to drug development and possible side effects of drugs to non-target biomarkers. Other methods such as genetic covariance (Shi *et al.*, 2016) can be used to identify genes shared across traits even where the causal variants differ.

Our method will not be able to identify genes where different causal variants have strong effects independently influencing each trait, and are in weak LD with each other. To account for these cases, we would need information on LD. However, we believe that the fact that *moloc* requires only summary association statistics and does not require LD estimates is advantageous, particularly when in-sample LD is not available, as misspecifications of LD data can lead to bias (Benner *et al.*, 2017). The statistics (priors and posteriors of configurations) will depend on the pattern of association (LD) and the number of SNPs in the region (Q) ((C. Giambartolomei *et al.*, 2014) and **Figure S10**). While *moloc* can be applied to any genomic region, complex loci such as the major histocompatibility complex (MHC) region, with extensive LD structure that exceeds the window size we consider in this analysis, would benefit from a tailored locus-based analysis (using genotyped information where possible).

Our goal is to find the functional relevance of genes to disease. This type of analysis differs to analysis using only GWAS to identify genes and pathways (Lamparter *et al.*, 2016). Although we identify a greater number of genes using SCZ GWASs only (Text S1), a joint analysis of multiple datasets provides additional information on the relevance of the functional information analyzed. We note that in our analysis, 53% of the genes we identified are novel, i.e. genes that fall outside of the PGC region. Future larger-scale GWAS would allow to confirm whether these novel associations are indeed true positives.

We expose one possible application of this approach in SCZ. In this application, we focus on scenarios involving eQTLs and GWAS, alone or in combination with mQTLs. While our method does not detect causal relationships among the associated traits, i.e. whether risk allele leads to changes in gene expression through methylation changes or vice versa, there is evidence supporting the notion that risk alleles might affect transcription factor binding and epigenome regulation that drives downstream alterations in gene expression (Tak and Farnham, 2015; Li *et al.*). We assign a prior probability that a SNP is associated with one trait (1 × 10^-4^), to two (1 × 10^−6^), and to three traits (1 × 10^−7^). We find support for our choice of priors in the data using two methods. The first uses stratified QQ plots (**Figure S4**). We find that eQTL enrichment in GWAS has a similar enrichment to mQTL in GWAS. The second is a form of empirical calibration as in Guo et al. (Guo *et al.*, 2015). We find that the prior and posterior expectations of colocalisation matched more closely under our choice of priors (**Table S5**). However the choices for prior beliefs for each hypothesis are always arguable. One could estimate priors for the different combinations of datasets. Pickrell et al. (Pickrell *et al.*, 2015) proposed estimation of enrichment parameters from genome-wide results maximizing a posteriori estimates for two traits. For multiple traits, another possibility is using deterministic approximation of posteriors (Wen *et al.*, 2017). We leave these explorations to future research. Additionally, instead of flat priors genome-wide, we can use priors that depend on per-SNP functional annotations. We provide the code and an example to do this using *fGWAS* (Text S1), and leave further applications to future research.

We note that this approach can be extended to more than three traits. Since the calculations are analytical and no recursive method is used, computation time for a region with 1000 SNPs is less than one second. However, time increases exponentially as number of traits increases. For four traits it is about 3 seconds, for five traits it is greater than 22 minutes. Overall, owing to the increasing availability of summary statistics from multiple datasets, the systematic application of this approach can provide clues into the molecular mechanisms underlying GWAS signals and how regulatory variants influence complex diseases.

## Acknowledgements

We would like to thank Chris Wallace at the Department of Medicine and MRC Biostatistics Unit, Cambridge Biomedical Campus, University of Cambridge, Cambridge, UK.

